# Graded and cross-reactive JAK–receptor interactions broaden the one-receptor, one-JAK rule

**DOI:** 10.64898/2026.10.24.761551

**Authors:** Mukesh Kumar Venkat Ramani, Philipp Ross, Kailin Guo, Yannik Leuz, Jillian Bonilla, Haley Hardtke, Juan L Mendoza

## Abstract

Cytokine receptors signal through a dedicated Janus kinase (JAK) that is thought to be constitutively bound to the receptor intracellular domain (ICD). We tested this one-receptor, one-JAK model biophysically. A bead-based screen of JAK1 and TYK2 against 32 receptor ICDs, confirmed by microscale thermophoresis to measure affinity, revealed dissociation constants spanning tens of nanomolar to tens of micromolar, indicating that many interactions are not high-affinity and may not be constitutive. Binding was also widely cross-reactive: GMCSFRα bound both JAK1 and TYK2, and IL2Rβ bound TYK2 far more tightly than JAK1. In cells, GM-CSF signaled through JAK1 when JAK2, JAK3, and TYK2 were inhibited. JAK–receptor pairing is therefore permissive and graded rather than exclusive and constitutive.

## Introduction

Cytokines, interferons and growth factors are secreted signaling proteins that coordinate hematopoiesis, immune responses and inflammation^1–4^. They act through transmembrane receptors that have no intrinsic catalytic activity^2^. Instead, each receptor chain depends on a cytoplasmic Janus kinase (JAK) for signal transduction. The JAK family comprises four non-receptor tyrosine kinases: JAK1, JAK2, JAK3, and TYK2^1–3^. Cytokine receptors fall into two structural classes. Class I (hematopoietin) receptors include the erythropoietin, growth hormone, and prolactin receptors, as well as shared-chain systems such as gp130 (IL-6 family), the common gamma chain (IL-2 family), and the common beta chain (IL-3 family). Class II receptors share a related but structurally distinct fold and are used by the interferons and the IL-10 family. For both classes, ligand binding dimerizes the receptor chains, bringing receptor-associated JAKs together intracellularly, so that they dimerize and cross-phosphorylate. The activated JAKs then phosphorylate the receptor and recruit STAT transcription factors^1–3^.

Receptor recognition is mediated by the JAK FERM and SH2-like domains, which fold together into a single integrated receptor-binding module^5,6^. This is distinct from the canonical functions of these domains in other proteins. The receptor intracellular domains (ICDs) are mostly intrinsically disordered and contain two short motifs, designated Box1 and Box2, that are critical for JAK binding. Box1 is a proline-rich, membrane-proximal segment that engages the F2 lobe and L1 linker of the FERM domain. It is the primary driver of binding affinity, and its deletion in receptors such as IFNλR1 abolishes stable JAK1 interaction^7^. Box2 is a hydrophobic segment 10–50 residues downstream, built around a conserved glutamate, that contacts the SH2 domain and stabilizes the complex when present. However, some ICDs lack a conserved Box1 and other ICDs are completely devoid of Box2^2,6,8,9^. While crystal structures of TYK2 bound to IFNαR1, JAK1 bound to IFNλR1 and IL10Rα, and a cryo-EM structure of full-length JAK1 bound to IFNλR1 highlight the structural basis for recognition^7,10–12^, structures of the JAK2 FERM–SH2 module bound to EPOR and the leptin receptor further show that receptor-mediated FERM dimerization is required for activation^13^.

JAK–receptor assignments were first established by co-immunoprecipitation and western blotting of endogenous or overexpressed proteins^14–17^. They are conventionally read as a one-receptor, one-JAK rule in which each receptor chain has a dedicated JAK partner. JAK2 serves the homodimeric receptors (EPOR, GHR, PRLR, TPOR) as a JAK2 homodimer, and JAK3 serves the common gamma chain. JAK1 serves IL2Rβ, IFNαR2 and IFNγR1, and TYK2 serves IFNαR1 and IL10Rβ^18,19^. Heterodimeric receptors therefore recruit two different JAKs, one per chain. The recognized exception is gp130, which associates with JAK1, JAK2 and TYK2^20,21^.

Underlying this paradigm is the premise that JAKs are constitutively bound to their receptors. In this view, each JAK is pre-associated with the Box1/Box2 region of its cognate ICD in the resting state, so that ligand-induced receptor rearrangement alone is sufficient to trigger activation^2,6^. JAK association also supports receptor surface expression^9^. This model implicitly requires two things. The first, constitutively bound JAK, implies the interaction to be stable, high-affinity binding between each JAK and its receptor. The second is that this binding is exclusive, so that each ICD is occupied by its assigned JAK and not by the others.

Neither requirement has been tested systematically across the receptor family with quantitative biophysics. Receptor–JAK assignments derive largely from cellular readouts such as co-precipitation and STAT phosphorylation. These integrate binding with expression levels, competition and downstream amplification, and so cannot report intrinsic affinity or reveal binding capacity that goes unused in cells. We therefore re-examine the paradigm on two fronts: First, that constitutive association predicts high-affinity binding for every receptor–JAK pair and second, that strict one-JAK-per-receptor pairing predicts selectivity. Structurally, however, Box1 and Box2 engage a conserved FERM–SH2 surface across JAK family members^6^, and receptor-mediated FERM dimerization is required for activation^6,13^. This suggests that any ICD capable of nucleating this dimeric geometry with a given JAK may be mechanistically primed to signal, whether that JAK is its canonical partner.

These questions matter clinically. Loss-of-function mutations in JAK3 or the common gamma chain cause severe combined immunodeficiency (SCID), and hypomorphic defects elsewhere in the pathway underlie a broader spectrum of primary immunodeficiencies^22,23^. Conversely, the somatic gain-of-function mutation JAK2 V617F drives myeloproliferative neoplasms, and the JAK family is implicated in rheumatoid arthritis, inflammatory bowel disease, psoriasis, and alopecia^24–29^. These diseases have prompted the development of many JAK inhibitors (JAKinibs)^30–37^. Several of them show cross-reactivity between JAK family members, and selective inhibition, especially of JAK1 versus JAK2, remains difficult^38–41^. This reflects in part the limited biochemical understanding of JAK–receptor interactions. If those interactions are more permissive and less uniformly constitutive than assumed, then which JAK engages which receptor in each cell may depend on context, with direct consequences for how inhibitors work.

Here we provide a systematic biophysical survey of JAK–ICD binding, using JAK1 and TYK2, two JAKs that undergo facile expression and purification, against a panel of 32 receptor ICDs. A bead-based screen identified binding events across receptor families, including many outside the reported canonical pairings. Affinity measurements by microscale thermophoresis (MST) confirmed these interactions and yielded affinity constants across a wide array of ICDs. Originally identified by the bead screen, we showed that one non-canonical pairing, GM-CSFRα with JAK1, supports signaling in cells when all other JAKs are rendered inactive using a combination of JAK inhibitors. Together, these data argue that JAK– receptor pairing is neither uniformly high-affinity nor exclusive, and they provide a framework for targeting JAK–receptor interactions more precisely.

## Results

### ICD sequence alignment shows indeterministic clustering for JAKs

To place the binding data in a structural context, we aligned the membrane-proximal Box1/Box2 region of each receptor ICD, grouped by their reported JAK partners (Fig. 1A). The sequence alignment highlights important prolines and multiple hydrophobic residues for Box1. The alignment highlights the diversity of information encoded through the ICD sequences. Apart from a few hydrophobic patches, the overall sequences show overall low conservation, even among ICDs reported to associate with the same JAK. However, this sequence divergence in the Box1/Box2 motifs further highlights the inherent diversity in signaling that can be achieved by altering the identity and affinity of the JAK-ICD interaction, which might provide for variations in kinetic activity of the JAKs.

**Figure 1.**
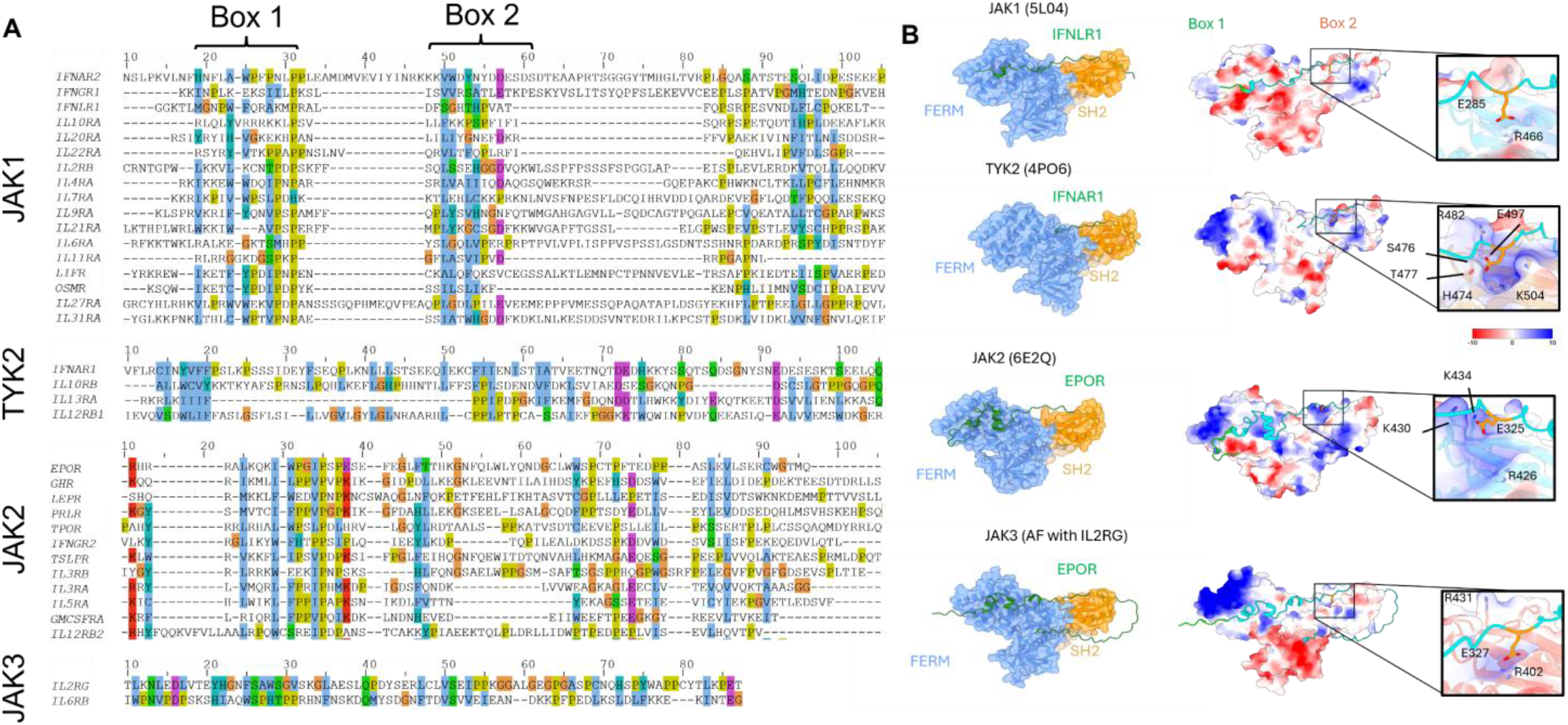
Sequence and structural analyses of JAK-ICD interactions. (A) Alignment of intracellular domains (ICDs) of cytokine receptors. Each alignment is done with ICDs of reported JAK partners. The coloring scheme utilized was Jalview’s Clustal scheme. (B) Structural interactions of JAKs with their reported ICDs, along with their electrostatic maps. The insets show the conserved glutamate of Box2 of the ICDs interacting with the SH2 domain of the JAKs. The JAK:ICD pairings are as follows - JAK1: IFNλR1 (PDB ID:5L04), TYK2: IFNαR1 (PDB ID: 4PO6), JAK2:EPOR (PDB ID: 6E2Q), JAK3: IL2Rg (AlphaFold Prediction). The FERM domains of the JAKs are colored in blue. The SH2 domains of the JAKs are colored in orange. The ICDs are colored green. Electrostatic maps are colored as indicated in the scale, with positive charges highlighted in blue and negative charges highlighted in red.

For structural context, we utilized the solved structures of JAKs bound to their canonical ICD partners: JAK1 – 5L04^11^, TYK2 – 4PO6^10^ and JAK2 – 6E2Q^13^ (Fig. 1B). Due to the unavailability of a solved structure for JAK3’s FERM-SH2 domains, we used AlphaFold structure prediction to generate a model of the JAK3 FERM-SH2 domains bound with the IL2Rγ ICD (Fig. 1B). Across all four JAK complexes, Box1 consists of a proline-rich motif sitting in a largely electronegative groove on the FERM domain. Box2 is more divergent among the receptors and engages a partially electropositive, hydrophobic patch spanning the FERM-SH2 junction. This asymmetry is consistent with Box1 setting the baseline docking energy for a FERM domain largely irrespective of which JAK it belongs to, while Box2 may fine-tune affinity^11^.

Most of the literature defines Box2 as a motif occurring around a conserved glutamate across all the JAKs. Empirical and predicted structures suggest the conserved glutamate is important for anchoring the ICD to the SH2-like domain of the JAKs by interacting with a partially electropositive pocket (Fig. 1B).

### Bead-based screening with JAK1 and TYK2 identifies different classes of receptor-JAK interactions

To get a comprehensive picture of JAK-ICD interactions, we purified SNAP-tagged JAK1 and TYK2 along with 32 receptor ICDs spanning the Box1 and Box2 motifs. JAK2 and JAK3 were not screened or measured because of limitations in protein expression and protein quality. We measured binding interactions across various families of ICDs with JAK1 and TYK2 (Fig. 2A). Initial observations of the data show a lower binding signal for JAK1 with canonical JAK1 interactors such as IFNγR1 and IFNαR2. One of the strongest known binders resulted in the highest fluorescence signal was IFNλR1 to JAK1, to which the entire dataset has been normalized (Fig. 2B). At first glance, the bead-based screen does not show clear binding preferences of ICDs to JAK1 or TYK2. However, our analysis can be used to group ICDs into multiple bins, or classes, based on their relative interaction with different JAK partners. For example, the entire IL-2 family of ICDs showed a higher binding signal to TYK2 than JAK1, whereas previous literature stated that all these ICDs interact with JAK1 but were assigned using whole-cell protein lysates and phospho-STAT signalling^42–47^. This type of cross-reactive interactions identified here may only be identified using binding assays and not using downstream signalling components as utilized in phospho-flow cytometry or western blotting.

**Figure 2.**
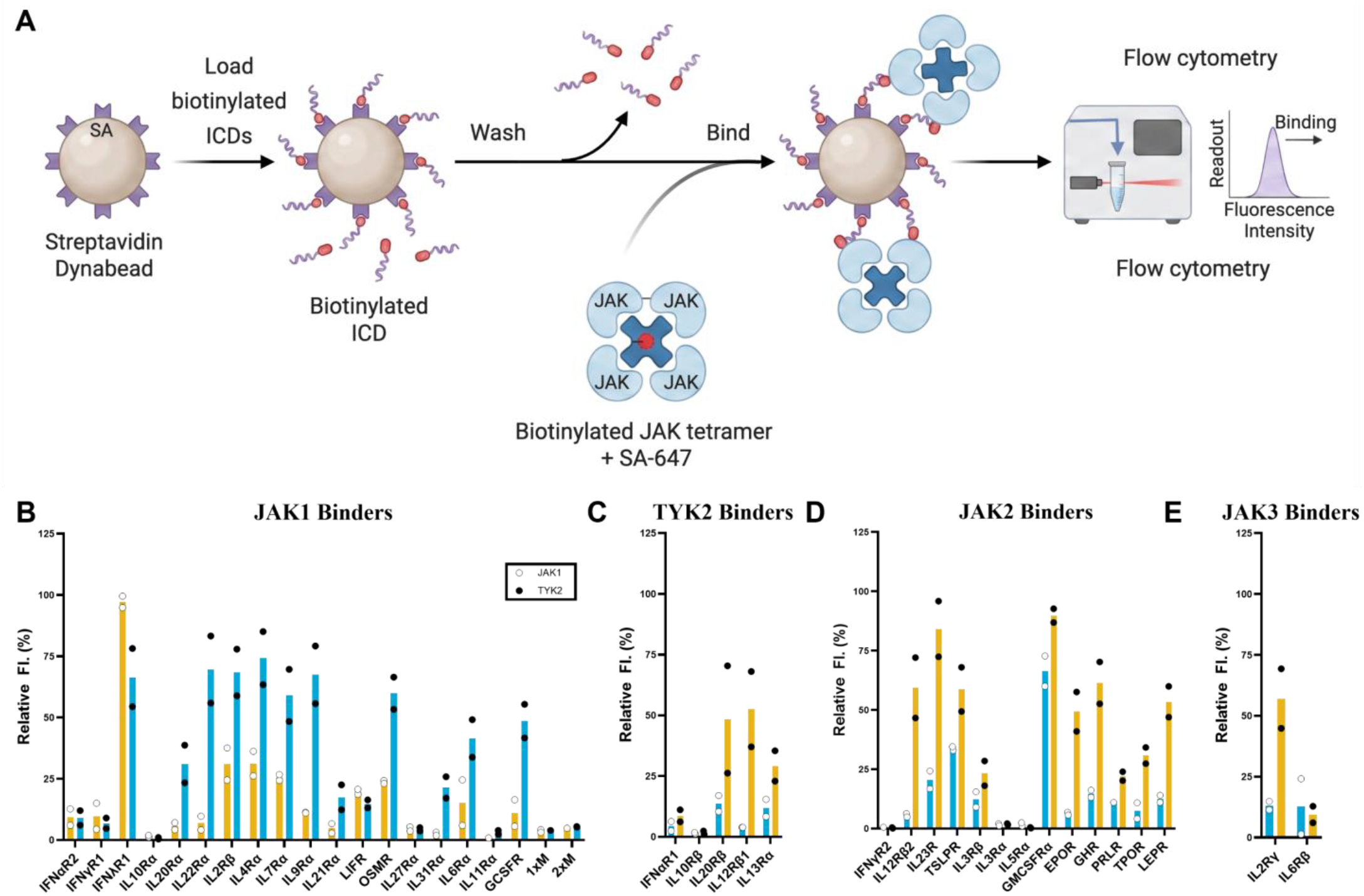
Bead screen assay showing binding interactions of ICDs with JAK1 and TYK2. (A) Workflow for bead screen assay using flow cytometry as the detection method. (B – E) ICD binding measured using the bead screen assay for reported JAK1 binders (B), TYK2 binders (C), JAK2 binders (D), and JAK3 binders (E). All experiments were duplicated with two technical replicates per biological replicate. Median Fluorescence values were normalized to the highest binding signal (JAK1’s interaction with IFNλR1).

### Confirmation of non-canonical and low-affinity interactions by microscale thermophoresis

To confirm binding trends identified in the bead screen by further using biophysical methods, we utilized a Monolith-X system to measure binding affinities by microscale thermophoresis (MST) coupled with spectral shift analysis. SNAP-JAK1 and SNAP-TYK2 constructs were labelled with the NanoTemper SNAP-Tag fluorophore kit and utilized for measuring affinities with 16 different purified ICDs (Table 1) (Fig. 3A). Our positive control for these experiments is JAK1 and IFNλR1, whose affinity has been reported as measured by surface plasmon resonance (SPR)^48^. Our affinity measurements show similar affinity values for JAK1 and IFNλR1, with a K_D_ of 381 nM (Fig. 3B). A new interaction for IFNλR1 that we measure here is with TYK2, with a K_D_ of 359 nM (Fig. 3C). This describes IFNλR1 as a receptor ICD that is both cross-reactive and does so with high affinity.

**Table 1.** Affinity measurements for JAK1 and TYK2 with selected ICDs of the IL-2 family, IL-10 family, and interferon family of receptor ICDs. All experiments were performed in triplicate; errors are ±SD.

| Receptor Families | ICD Names | JAK1 | TYK2 |
| --- | --- | --- | --- |
|  |  | K<sub>D</sub> (μM) | K<sub>D</sub> (μM) |
| Interferons | IFNαR1 | 3.09 ± 1.23 | 0.048 ± 0.04 |
|  | IFNαR2 | 2.93 ± 2.85 | 5.78 ± 9.51 |
|  | IFNγR1 | 5.16 ± 2.53 | 9.74 ± 6.98 |
|  | IFNγR2 | 4.85 ± 2.11 | 7.01 ± 4.17 |
|  | IFNλR1 | 0.381 ± 0.13 | 0.359 ± 0.30 |
| IL-10 Family | IL10Rβ | 7.11 ± 4.61 | 10.9 ± 7.63 |
|  | IL20Rα | 12.4 ± 5.25 | 18.9 ± 11.70 |
|  | IL22Rα | 19.6 ± 15.21 | 13.7 ± 3.51 |
| IL-3 Family | GMCSFRα (CSF2Rα) | 0.785 ± 0.36 | 1.34 ± 0.85 |
|  | IL3Rβ | 29.6 ± 6.69 | 2.69 ± 3.30 |
| Control | 2xMUT LR1 | 2.23 ± 1.01 | Not measurable |
| IL-2 Family | IL2Rβ | 3.95 ± 2.34 | 0.068 ± 0.09 |
|  | IL2Rγ | 6.71 ± 4.08 | 9.94 ± 3.75 |
|  | IL4Rα | 7.69 ± 4.68 | 2.09 ± 1.00 |
|  | IL7Rα | 31.4 ± 10.45 | 44 ± 18.76 |
|  | IL9Rα | 19.2 ± 8.25 | Not measurable |

**Figure 3.**
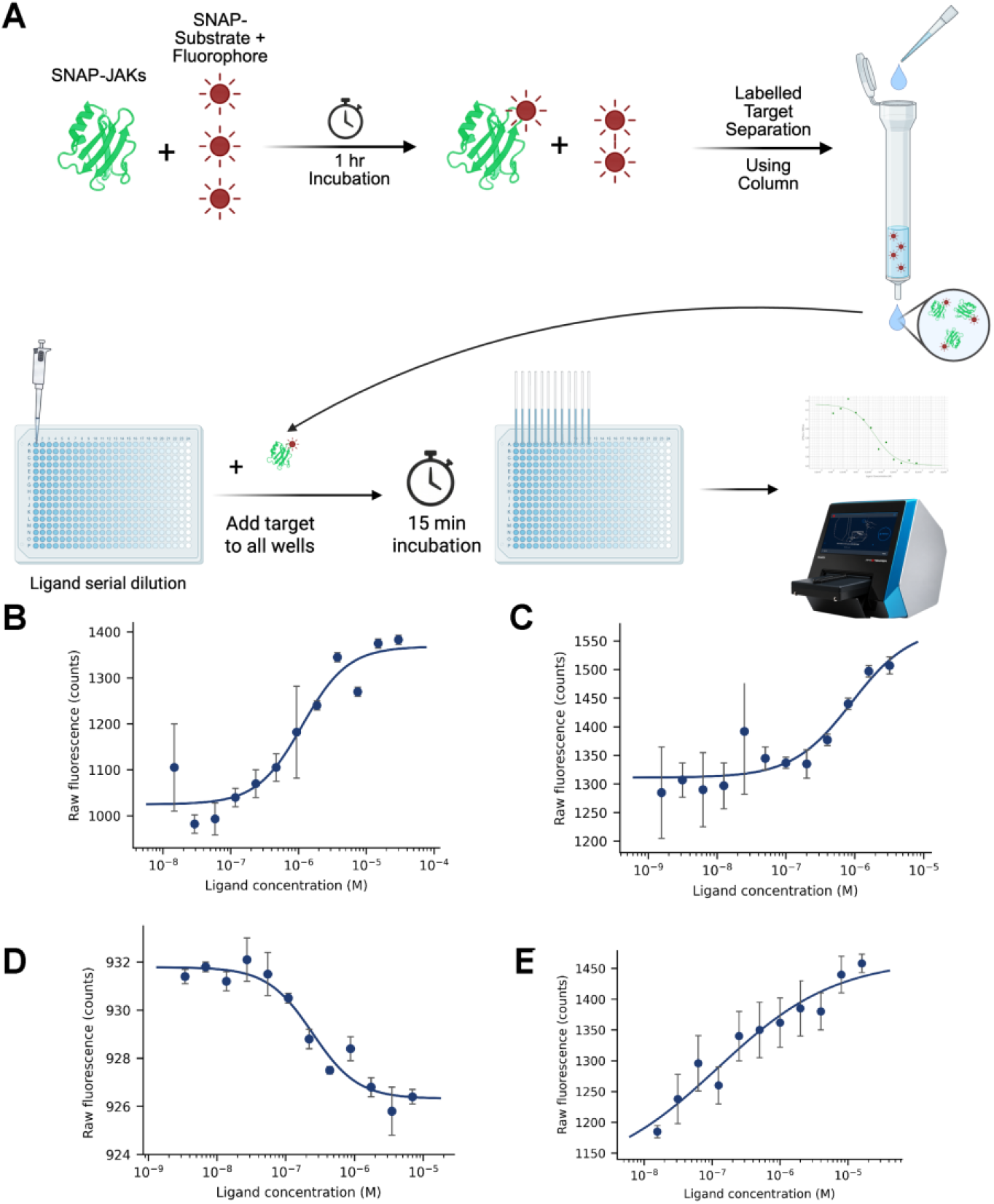
Determination of K_D_ for IFNλR1 and GMCSFRα by MST. (A) Workflow for Monolith SNAP-Labeling and binding assays. (B – E) Binding curves for JAK1 with IFNλR1 (B), TYK2 with IFNλR1 (C), JAK1 with GMCSFRα (D), and TYK2 with GMCSFRα (E). Experiments were performed in triplicate; data were fitted with a K_D_ model (1:1 binding), with error shown as ±SD.

The bead screen identified an additional high-affinity and cross-reactive receptor ICD. GMCSFRα was found to interact with both JAK1 and TYK2 with a very high signal in the bead screen (Fig. 2B). GMCSFRα has a short 58-amino-acid ICD which is extremely important for JAK2 binding and signal transduction, as evidenced by deletion studies^49^. The GMCSF-recruited complex is described as only dependent on JAK2 for signaling. Therefore, we aimed to validate GMCSFRα binding to JAK1 and TYK2 as measured in the bead screen. For JAK1, GMCSFRα was measured to have a K_D_ of 758 nM (Fig. 3D), and similarly, a K_D_ of 1.34 µM for TYK2 with GMCSFRα (Fig. 3E). Like IFNλR1, GMCSFRα supports receptor-ICD interactions that can be cross-reactive but bind with high affinity to different JAKs.

### GMCSFRα’s use of a non-canonical JAK interaction supports signaling in cells

Biophysical data demonstrate that GMCSFRα’s ICD can bind to JAK1 and TYK2, two predicted non-canonical interactions. However, this raises the question of whether the binding of GMCSFRα to JAK1 or TYK2 is capable and sufficient to support signaling in a cell when no other JAKs are active. To measure signaling, we utilized available JAK inhibitors (JAKinibs) in different combinations to achieve single or two JAKs active at a time to compare with the native JAK2-mediated signaling (Fig. 4A). In these experiments, we show via a phospho-STAT5 assay that GM-CSF signals strongly, 67.6% compared to GM-CSF without inhibitors, when only JAK1 is active (Fig. 4B). By contrast, when only JAK2 is active, the canonical paired cytokine to GMCFSRα, GM-CSF signals about 23.3% compared to uninhibited GM-CSF. When only

**Figure 4.**
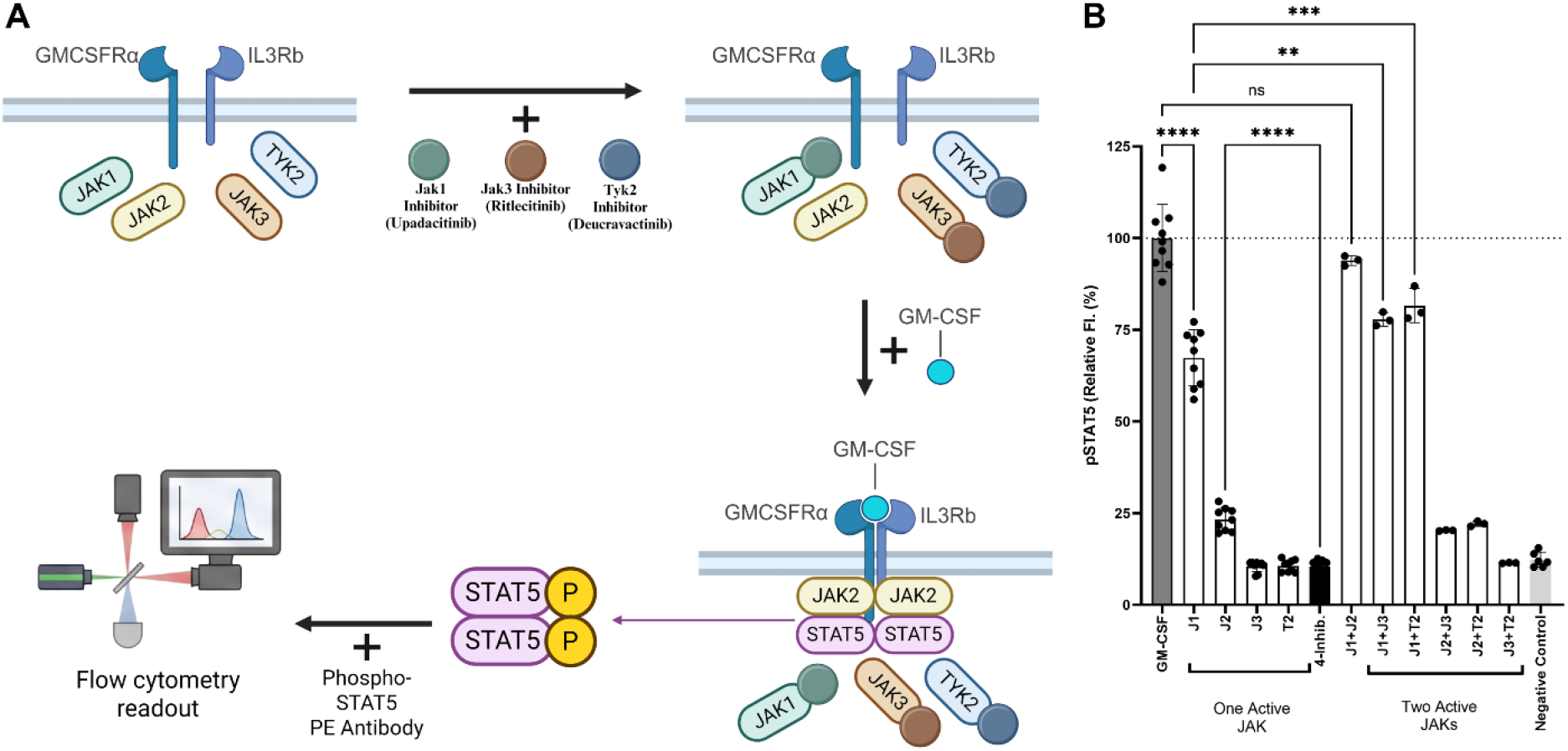
pSTAT5 signaling assay using JAK inhibitors demonstrates the contributions of non-canonical JAKs to signaling. (A) Workflow for phospho-flow assay using flow cytometry. (B) pSTAT5 fluorescence induced by GM-CSF signaling using JAK inhibitors. J1 = JAK1, J2 = JAK2, J3 = JAK3 and T2 = TYK2. For each active JAK sample, multiple inhibitors were added to inhibit the non-active JAKs for that sample. (For the J1 active sample, inhibitors for J2, J3, and T2 were added). Experiments were performed in triplicate for the double inhibitor study, and n=9 was used for the single inhibitor study. Significance was established using pairwise ANOVA, ns = p>0.05, * = p<0.05, ** = p<0.01, *** = p<0.001, **** = p<0.0001.

JAK3 or TYK2 is active, we did not detect a measurable signal compared with the untreated sample. To further test how multiple JAKs, both canonical and non-canonical, contribute to signaling, we used combinations of two JAKinibs so that pairs of JAKs would remain active. When both JAK1 and JAK2 were active, such that JAKinibs for TYK2 and JAK3 were used, cells signaled at levels equal to those of GM-CSF-treated samples without any inhibitors. Interestingly, when JAK1 was active in combination with JAK3 or TYK2, cell signaling showed a higher signal compared to only JAK1 being active.

## Discussion

The one-receptor, one-JAK model rests on two assumptions: that each receptor chain is stably occupied by a dedicated kinase, and that this pairing is exclusive. Our biophysical survey of JAK1 and TYK2 against 32 receptor intracellular domains (ICDs) supports neither. Affinities spanned nearly three orders of magnitude, from 48 nM (TYK2–IFNαR1) to tens of micromolar, and several canonical pairs were weak: IFNαR2 and IFNγR1 bound JAK1 with dissociation constants of 3–5 μM. Binding was also frequently promiscuous. TYK2 bound the IL2Rβ ICD with an affinity (68 nM) comparable to that for its canonical partner IFNαR1 and more than 20-fold tighter than JAK1, and the GM-CSFRα ICD bound both JAK1 and TYK2 (0.8 and 1.3 μM). This interaction is functional: when JAK2, JAK3, and TYK2 were inhibited, GM-CSF-induced STAT5 phosphorylation retained about 65% of its uninhibited level through JAK1. Binding of JAK1 to the common β chain has been reported (51); our measurements quantify it and show that it extends to a receptor conventionally assigned to JAK2.

These data suggest that JAK–receptor association is better described as a graded, competitive equilibrium than as constitutive wiring. The tightest pairs are likely constitutive: IFNαR1 and EPOR depend on TYK2 and JAK2, respectively, for cell-surface expression (52, 53). But interactions in the micromolar range are unlikely to be saturated at cellular JAK abundance unless membrane confinement or receptor oligomerization raises the effective concentration. Ligand-induced receptor dimerization, which creates an additional receptor contact and stabilizes a JAK2 FERM dimer (13), may therefore not only activate JAKs but also recruit or retain them, converting weak, transient interactions into productive complexes.

Occupancy alone does not determine output, however. A receptor-bound JAK signals only if it pairs productively with a partner, and pairs probably differ in efficiency because dimer geometry, pseudokinase-mediated dimerization and trans-phosphorylation depend on the identity of both kinases (12, 13). Our phospho-flow data are consistent with this. At the GM-CSF receptor, JAK1 alone supported about 65% of the uninhibited STAT5 signal, whereas JAK2 alone, the canonical partner, supported about 25%. JAK1 with JAK2 restored the full signal, and JAK1 with JAK3 or TYK2 exceeded JAK1 alone. Pair-specific coactivation efficiency could account for this ranking. So could unequal JAK1 and JAK2 abundance in THP-1 cells, residual inhibition of JAK2 by the JAK1-directed compound, or a difference in JAK2 affinity for the receptor, which we did not measure. These explanations can be separated by genetic deletion, measurement of JAK levels, and JAK2–ICD affinities.

Together, these layers should determine how readily one JAK can rescue another: rescue potential should scale with the occupancy of the ICD by the alternative kinase and with the activation efficiency of the pair it forms. A modest binder can therefore dominate where it is abundant, whereas a tight binder expressed at low levels is a poor backup, and the same receptor may be rescuable in one lineage but not another. This prediction is compatible with pharmacological data. In myeloproliferative neoplasms, cells persisting under JAK2 inhibition reactivated signaling through heterodimers of JAK2 with JAK1 or TYK2 (54). In patients with rheumatoid arthritis receiving tofacitinib, cytokine-induced STAT phosphorylation was reduced by only 10–73%, depending on cytokine and cell type, with the weakest effect on IL-10–induced STAT3 phosphorylation in monocytes (55). Pharmacokinetic–pharmacodynamic modeling at approved doses likewise predicts incomplete daily inhibition of several pathways, including IL-4 and IL-2/IL-15 (56). Residual signaling is usually attributed to drug exposure and kinase selectivity, and our data does not exclude this. They do identify receptors in which an alternative JAK is poised to contribute; the IL4Rα ICD, for example, bound TYK2 more tightly than JAK1. Conversely, the allosteric TYK2 inhibitor deucravacitinib blocks IL-12, IL-23 and type I interferon signaling but spares JAK1/JAK3-dependent cytokines (57), although TYK2 binds the IL2Rβ ICD with high affinity in vitro. Binding capacity therefore need not imply functional requirement; it may define a latent reserve that matters only when the primary JAK is lost or blocked.

If so, selectivity engineered at the ATP site must contend with redundancy at the receptor. Disrupting the FERM– SH2/ICD interface, or combining JAK-selective agents, may be required to silence pathways that can recruit an alternative kinase.

Our study does have limits, as we tested only JAK1 and TYK2; JAK2 and JAK3 were not screened or measured because of limitations in protein expression and protein quality. We also used isolated FERM–SH2 modules and ICD fragments outside a membrane, so avidity, pseudo kinase contributions, and receptor dimerization are absent. Either one or a combination of factors may affect signaling even though there are confirmed and measurable receptor-JAK affinities. The occupancy relation assumes one binding site per ICD and equilibrium with unknown free JAK concentrations. The cellular assay relies on small-molecule inhibitors whose selectivity is not absolute. Extending the survey to JAK2 and JAK3 and measuring JAK expression and occupancy of intact receptors in genetically defined cells, will test the model directly. JAK–receptor specificity is better viewed as a tunable, competitive landscape than as a fixed wiring diagram, and cytokine biology and JAK-targeted therapy will need to account for it.

## Methods

### Sequence alignments

Sequences were truncated at the transmembrane domain and aligned using default MUSCLE parameters in Jalview. Alignment was cropped at N and C terminus to emphasize regions of conservation, and residues were colored using the Clustal scheme implementation in Jalview. JAKs were truncated to FERM-SH2 regions using alignment as reported^50^. Electrostatic map was calculated in ChimeraX.

### Cloning and protein purification

Protein constructs were ordered as twist fragments and cloned into pDG vectors using Gibson cloning. The cloned vectors were confirmed using Sanger sequencing and transformed into NEB Express Iq Competent *E. coli* (NEB: C3037I) for protein expression. Transformations were carried out by thawing cells on ice for 10 min, adding 50 ng of DNA and incubating on ice for 10 min, heat-shocking at 42 °C for 75 seconds, and cooling the cells on ice for 5 min. The cells were recovered in SOC medium for 1 h at 37 °C and were plated on Luria−Bertani agar plates with 50 μg/mL kanamycin for selection using the spread plate technique. Individual colonies were grown in Luria-Bertani medium at 37 °C containing 50 μg/mL kanamycin in 100 mL flasks. 100ml of inoculum was used for inoculating 1 L of terrific broth (Thermo Fisher), and the culture was grown for 1 hour at 16 °C. IPTG was used to induce the expression at a final concentration of 0.5 mM. The cultures were pelleted by centrifugation after overnight growth (16 h at 16 °C), and the cells were lysed by sonication in a lysis buffer (50 mM HEPES-NaOH pH 7.5, 500 mM NaCl, 10% glycerol, 0.1% Triton X-100, 15 mM imidazole, 10 mM BME, 1 mg/ml DNase I and 2 tablets of Protease Inhibitors (Millipore – 11836170001)). The sample was homogenized in an Avestin C5 emulsifier at 10000-15000 psi for 3 passes. The lysate was clarified by centrifugation at 50000 g for 30 min at 4 °C. The proteins were purified through affinity column chromatography using Ni2+/NTA beads (Qiagen). Briefly, the column was equilibrated with lysis buffer, and the cleared lysate supernatant was run through the column. A wash (50 mM HEPES-NaOH pH 7.5, 500 mM NaCl, 10% Glycerol, 25 mM Imidazole and 10 mM BME) was done before eluting with an elution buffer (20 mM HEPES-NaOH pH 7.5, 500 mM NaCl, 10% Glycerol, 250 mM imidazole, and 10 mM BME). Proteins were concentrated using centrifugal filtration and cleaned up by size exclusion chromatography using a Superdex 200 column (Cytiva). The integrity of the proteins was assessed by polyacrylamide gel electrophoresis.

### A bead-based screening method of cytokine receptor interactions with JAK1 and TYK2

To assess binding of JAKs to the ICDs, bead-immobilized biotinylated-ICD targets were generated using a streptavidin capture adapted from the manufacturer’s recommendations. Streptavidin-coated Dynabeads (Dynabeads M-280, Thermo Fisher Catalog No. - 11205D) were equilibrated by one wash in bead buffer (20mM HEPES pH 7.5, 300mM NaCl, 0.1% BSA, 0.01% Tween-20) using a magnetic particle separator (Miltenyi Biotec MACS Separator) and resuspended in their original volume.

For each ICD construct to be immobilized, 20 µl of washed bead slurry was combined with 400 pmol of biotinylated ICD protein and brought to a final volume of 30 µl with bead buffer (10-fold excess over specified binding capacity). The reactions were incubated overnight at room temperature in the dark, followed by an additional 1 hour at 4 °C with periodic mixing. ICD-loaded beads were washed four times with 80 µl bead buffer using magnetic separation, resuspended in a final volume of 80 µl of bead buffer, and stored at 4 °C for a maximum of 1 month.

For staining, biotinylated JAK FERM-SH2 constructs were tetramerized by direct mixing with fluorophore-conjugated streptavidin (SA-AF647) to a final concentration of 100 nM and incubated for 30 mins at room temperature in the dark. Free SA-AF647 was quenched by addition of freshly prepared D-biotin (adjusted to a pH of 7) to a final concentration of 500 nM for 5 mins, after which tetramers were diluted to a working concentration in bead buffer and kept at 4°C until use. Immediately before staining, ICD-loaded bead suspensions were thoroughly homogenized by pipetting to ensure uniform bead distribution, and 1 µl of slurry was combined with 17 µl of JAK FERM-SH2 tetramer solution in a U-bottom 96-well plate (Two technical replicates per ICD construct). Staining reactions were incubated for 30 min at room temperature in the dark. Beads were pelleted by centrifugation (3500 × g for 3 min at 4 °C), washed once with 200 µl of bead buffer before being resuspended in 200 µl of bead buffer. Samples were analyzed by flow cytometry.

### SNAP-labeling for microscale thermophoresis measurements

SNAP-JAK1 and SNAP-TYK2 were labelled with SNAP-TAG labelling kits from NanoTemper (MO-L019 – Red labelling). 150 µl of 15 µM protein was supplemented with 1 mM DTT and utilized for labelling by adding 10 µl of 200 µM dye (Per manufacturer’s instructions). The incubation was carried out for 1 hour at room temperature. Unlabeled dye was separated using a manufacturer-provided column from the kit. Efficiency of labelling was calculated by utilizing degree of labelling calculations per manufacturer’s instructions. The degree of labelling for SNAP-JAK1 was estimated at 0.51, and the same for SNAP-TYK2 was 0.58.

For monolith experiments, 10 µl of 40 nM labelled protein (Target) was mixed with 10 µl of purified ICDs (Ligand) (Serially diluted into 12 wells) in a 384-well plate. All initial ligand concentrations are highlighted in Table 1. The reaction was incubated for 15 minutes and aspirated into premium capillaries (NanoTemper - MO-K025). The capillaries were assayed for fluorescence signals at 650nm and 670nm using a Monolith X system (NanoTemper).

### Signaling assay

THP1 cells were cultured and seeded at 0.02 million cells per well in a 96-well plate. JAK inhibitors were added at concentrations which are 10 times more than recommended IC50 times as a guiding principle for inhibition. Upadacitinib (JAK1 inhibitor) was used at 100 nM wherever JAK2 is not inhibited and at 1000 nM when JAK2 is also inhibited. Fedratinib (JAK2 inhibitor) was used at 30 nM. Ritlectinib (JAK3 inhibitor) was used at 500 nM. Deucravactinib (TYK2 inhibitor) was used at 100 nM. All cells were incubated with inhibitors for 30 minutes. The cells were induced with 5 nM of GM-CSF ligand (MedChem - HY-P7016A). After incubation for 15 minutes with GM-CSF ligand, cells were fixed in 2% PFA for 10 mins at room temperature and spun down at 1500g for 4 mins. The supernatant was discarded, and the cells were permeabilized using 100% methanol for 1 hour on ice. After washing 3 times with PBS + 0.5% BSA, cells were stained with Phospho-STAT5 PE conjugate (BioLegend – 936904) at a 1:1000 dilution for 1 hour. After washing 3 times with PBS + 0.5% BSA, fixed cells were analyzed using flow cytometry.

## Acknowledgements

Funding to support J.L.M. is provided by the Pritzker School of Molecular Engineering at the University of Chicago, the National Institutes of Health (NIH) grant 1R35GM147179-01, and the Howard Hughes Medical Institute.

## Author contributions

All authors contributed to writing and reviewing the manuscript. J.L.M. conceived of the project, contributed to the design of the studies, and oversaw experiments. M.K.V.R and P.R. contributed to overall experimental design and cloning. M.K.V.R carried out the Monolith binding measurements and signaling assays. P.R. carried out the bead-based screening assay and optimized Monolith binding measurements. K.G. did the sequence alignments and structural prediction with AlphaFold. Y.L., J.B. and H.H. helped with protein purification of the JAK and ICD constructs.

